# Oligodendrocyte precursor cells establish contacts with somata of active neurons

**DOI:** 10.1101/2025.03.28.646001

**Authors:** Ying Sun, Peter Jukkola, Yetunde Akinlaja, Olga Garaschuk, Friederike Pfeiffer

## Abstract

Oligodendrocyte precursor cells (OPCs) generate myelinating oligodendrocytes and interact with synapses to regulate neuronal networks. Neuronal activity induces proliferation or differentiation of OPCs, but it is unknown whether OPCs in turn affect neuronal activity. Here, we show that OPCs establish close contacts with the somata of active neurons. We found that neurons contacted by OPCs and their processes display reduced calcium signals, indicating that they regulate their activity.

## Main Text

Oligodendrocyte precursor cells (OPCs) are mainly known for generating oligodendrocytes (OLs), essential for proper myelination in the central nervous system (CNS). Recently revealed additional functions of OPCs include remodeling of growing axons ^1^ and synapses ^2,3^, or facilitating exocytosis of neuronal lysosomes ^4^. These findings support the concept of OPCs exerting functions beyond the generation of OLs. OPCs receive synaptic inputs from neurons ^5,6^ and are able to sense neuronal activity^7^, which can induce their proliferation or differentiation into mature oligodendrocytes ^8^. However, it is not clear whether this interaction is bidirectional.

In white matter, OPCs have been shown to contact neurons with their numerous processes at discrete locations such as the nodes of Ranvier ^9,10^. Therefore, we first explored spatial relationships between OPC processes and neuronal subdomains in the gray matter.

PDGFRα immunolabeling of a postnatal day 14 (P14) hippocampus showed OPC cell bodies in strata oriens and radiatum, where axons and dendrites localize, but some were located at the border and occasionally within the pyramidal layer, where the somata of pyramidal neurons are arranged (Extended Data Fig. 1). Highly branched processes of OPCs covered the neuropil, and some extended across the stratum pyramidale (Fig. 1a), surrounding the somata of pyramidal neurons labeled with microtubule-associated protein 2 (MAP-2) (Fig. 1a, b). Other OPC processes intertwined with MAP-2-positive apical and basal dendrites of neurons (Fig. 1a, c-e) and the axon initial segments (AIS) identified by βIV-spectrin (Fig. 1a, d). In adult mice, we found the same close association between OPC processes and the somata of cortical layer 5 pyramidal neurons labeled with Kv2.1, a commonly accepted marker for neuronal somata (Fig. 1f) and the AIS of cortical pyramidal neurons (Fig. 1f-h). Thus, OPC processes approach soma, dendrites and AIS of neurons in the hippocampus of juvenile (Fig. 1a-e) and adult mice (Fig. 1f-h), in addition to their well-described interactions with axons prior to myelination.

**Figure 1.**
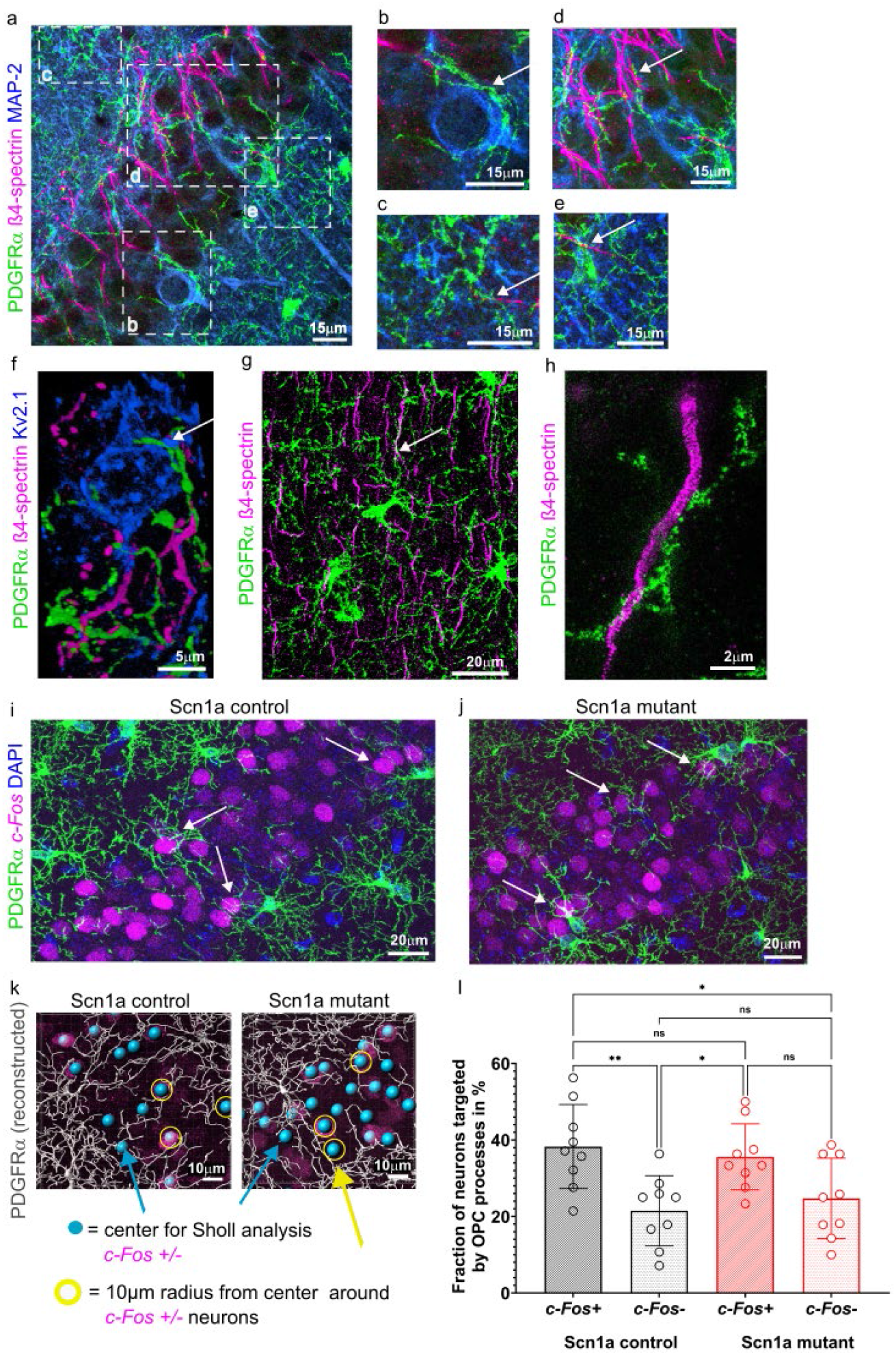
OPCs target active neuronal somata in hippocampus and cortex of juvenile and adult mice. a-e. Immunolabeling for PDGFRα showing the distribution of OPCs in stratum oriens (so), stratum pyramidale (sp) and stratum radiatum (sr) of the hippocampal CA1 region at P15, where they approach various neuronal structures with their protrusions (white arrows). Neuronal structures are stained with anti-ßIV spectrin (axon initial segment (AIS); magenta) and anti-MAP-2 (soma and dendrites; blue) antibodies. Scale bars: 15 µm. b-e show zoomed in areas indicated by white boxes in a. f. OPC processes stained with anti-PDGFRα antibody (green) around the soma of a layer 5 cortical pyramidal neuron expressing Kv2.1 (blue) and the AIS (stained with anti-ß4-spectrin antibody; magenta). Scale bar: 10µm. g. OPCs (stained with anti-PDGFRα antibody; green) interacting with neurons and their AIS (stained with anti-ß4-spectrin antibody; magenta) in layer 2-3 of the cortex. Scale bar: 20µm. h. high magnification (STED microscopy) showing OPC processes (stained with anti-PDGFRα antibody; green) twisted around the AIS (stained with anti-ß4-spectrin; magenta) of a cortical pyramidal neuron. Scale bar: 5µm. i-j. Confocal images of CA1 in Scn1a control (i) and mutant (j) mice showing OPC processes (stained with anti-PDGFRα antibody; green) and active neuronal somata (stained with anti-*c-Fos* antibody; magenta) in the stratum pyramidale. Nuclei are labeled with DAPI (blue). Scale bars: 20 µm. k. Design of Sholl analysis from 3D reconstructions. Sholl analysis was performed by positioning the starting point for automated filament reconstruction in Imaris onto either *c-Fos*+ (red), or *c-Fos*-(grey) neuronal soma in CA1 stratum pyramidale. And l. The number of neurons that were surrounded by 5-10 OPC processes within 10-µm distance around the soma center of the neuron was assessed per image. 5 images per mouse (n=3) were analyzed. Kruskal-Wallis test with Dunn’s multiple comparison test was performed (control *c-Fos*+ vs. control *c-Fos*-p=0.0035; control *c-Fos*+ vs. mutant *c-Fos*+ p>0.9999; control *c-Fos*+ vs. mutant *c-Fos*-p=0.0421; control *c-Fos*-vs. mutant *c-Fos*+ p=0.0038; control *c-Fos*-vs. mutant *c-Fos*-p>0.9999; mutant *c-Fos*+ vs. mutant *c-Fos*-p=0.0450).

To examine how the spatial relationship between OPCs and neuronal somata changes upon an increase in neuronal activity in neonate mice, we used P14 *Vgat*^*cre/+*^*;Scn1a*^*A1783V/+*^ mice (Scn1a mutant) as a model of genetic epilepsy (Kuo et al., 2019; Ricobaraza et al., 2019; Extended Data Figure 2).

There was no difference in the density of PDGFRα-immunolabeled OPCs in Scn1a control and mutant CA1 (Extended Data Fig 3) and we could not detect major changes in overall OPC morphology in Scn1a mutant mice by performing 3D Sholl analysis (Extended Data Fig. 4a), except for a slight increase in the stratum oriens in specific intersections within 18-27µm away from the soma center when we separated OPCs by location (Extended Data Fig. 4b). There were no changes in the number of intersections between mutant and control OPCs localized in the stratum pyramidale and radiatum (Extended Data Fig. 4c-d). In Scn1a+/-heterozygous mutants, parvalbumin-expressing interneurons show normal firing through postnatal day 10 (P10) followed by a sudden onset of reduced firing rate at P11 ^11^. Thus, the relatively short duration of the mildly hyperexcitable state may not have been sufficient to elicit a proliferative response (Extended Data Figure 3) or major changes in OPC morphology (Extended Data Figure 4).

We next determined whether OPC processes were more prevalent around active neurons expressing the immediate early gene product *c-Fos*, whose transcriptional activity has been used as an indicator of actively firing neurons ^12^. In both control and mutant mice, *c-Fos+* neurons were surrounded by PDGFRα-positive OPC processes (Fig. 1i-j). The percentage of neurons that were surrounded by 5-10 OPC processes within 10-µm distance around the soma center of the neuron (Fig. 1k-l) was assessed by Sholl analysis, placing the center of the Sholl spheres onto either *c-Fos+* or *c-Fos-*neurons (Fig. 1k). In control mice, there were significantly more *c-Fos+* neurons that were contacted by 5-10 OPC processes emanating from nearby OPCs within a 10µm radius as compared to *c-Fos-*neurons (Fig. 1l). There was no significant difference in the number of *c-Fos*+ neurons targeted by OPC processes between control and mutant mice, nor was there a significant difference between *c-Fos-*neurons. Thus, under physiological conditions, OPC processes preferentially contacted nearby *c-Fos+* active neurons, suggesting that neuronal activity is a key determinant for attracting OPC processes towards them.

Since the short lifespan of the Scn1a mutant mice precluded analysis beyond P14, we used a chemogenetic approach to activate pyramidal neurons in adult mice by using Camk2a-cre (T29-1) tg/+; CAG-LSL-Gq-DREADD (CAG-CHRM3-mCitrine) tg/+ double transgenic mice. In dcz (deschloroclozapine, to induce the canonical Gq pathway) - injected mice, *c-Fos* expression was prominently elevated in CA1 compared with *c-Fos* in vehicle-injected mice (Figure 2a, b), in line with previous reports ^13^. With the dcz-injected animals showing a clear increase in *c-Fos+* neurons, we assessed the interaction between OPCs and their processes and *c-Fos*+ neuronal somata in the CA1 region with regard to the fluorescence intensity of *c-Fos* (Figure 2c-d). In control vehicle-injected mice, the fluorescence intensity of *c-Fos* was significantly lower in neuronal somata contacted by OPC processes as compared to those without (Figure 2e, supplementary movie 1). The same pattern occurred in dcz-injected DREADD mice (Figure 2f, supplementary movie 2).

**Figure 2.**
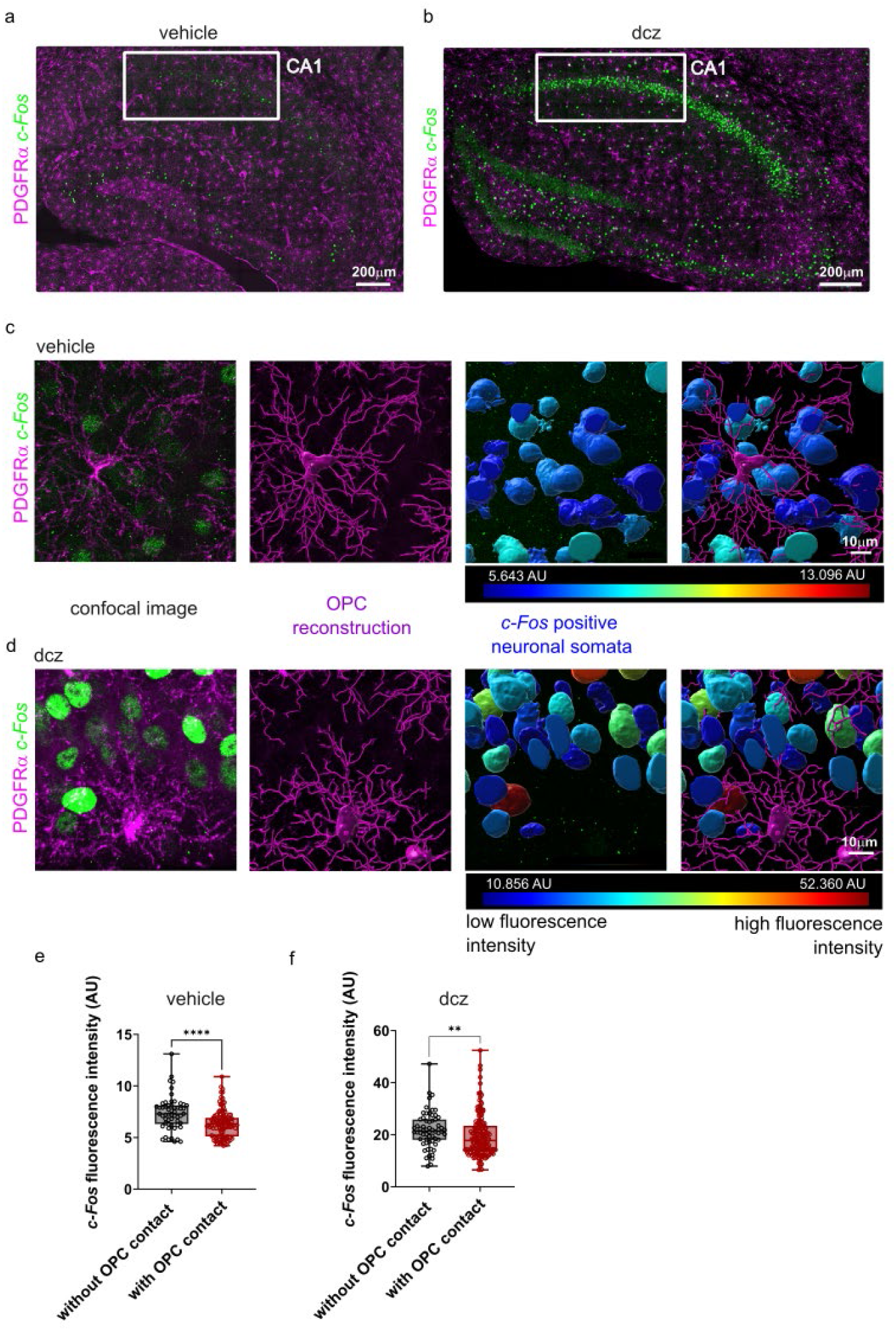
Neurons targeted by OPCs in the adult hippocampal CA1 region show weaker fluorescence intensity of *c-Fos* staining. a-b. *C-Fos* (green) expression 6 hours after vehicle (a) or dcz (b) injection in the hippocampus of Camk2a-cre (T29-1) tg/+; CAG-LSL-Gq-DREADD (CAG-CHRM3-mCitrine) tg/+ adult mice. OPCs are labeled with anti-PDGFRα antibody (in magenta). White boxes show CA1 and CA2/3 regions, respectively. Scale bars: 200µm. c. vehicle: OPC (PDGFRα-labeled, magenta) localized in the pyramidal layer of CA1 in between *c-Fos*+ neurons (green). OPC soma were reconstructed using the surface tool (magenta) and filaments were reconstructed using the filaments tool (magenta) in Imaris. *C-Fos*+ neurons were reconstructed with the surface tool (color-coded for intensity). Scale bar: 10µm. d. dcz: OPC (PDGFRα-labeled, magenta) localized in the pyramidal layer of CA1 in between *c-Fos*+ neurons (green). OPC soma were reconstructed using the surface tool (magenta) and filaments were reconstructed using the filaments tool (magenta) in Imaris. *C-Fos*+ neurons were reconstructed with the surface tool (color-coded for intensity). Scale bar: 10µm. e. Assessment of differences in *c-Fos*+ fluorescence intensity between neuronal somata in the CA1 region with or without OPC contact in the vehicle-injected DREADD mouse. n=174 cells, Mann-Whitney test (two-tailed) p<0.0001 (sum of ranks 6096, 9130). f. Assessment of differences in *c-Fos*+ fluorescence intensity between neuronal somata in the CA1 region with or without OPC contact in the dcz-injected DREADD mouse. n=236 cells, Mann-Whitney test (two-tailed) p=0.0033 (sum of ranks 8696, 19270).

Our observations indicate that OPCs extend their processes towards active neurons altering the level of their *c-Fos* expression. To assess the direct physiological impact of this spatial relationship, we measured changes in calcium signals reflecting spontaneous activity in pyramidal neurons of the hippocampus in neonate mice ^14^. In acute slices of neonate mice expressing the fluorescent protein DsRed under the control of the NG2 promotor (Fig. 3a,b), the average change in fluorescence was compared between neurons directly in contact with OPCs (identified by expression of DsRed) and neurons without OPC contact (Fig. 3a-c). Neurons in contact with OPCs displayed significantly lower changes in calcium levels as compared to those without apparent OPC contact (Fig. 3d), indicating that OPCs may influence neuronal activity by surrounding the soma with their processes. Intensity based reconstruction of neuronal somata based on their *c-Fos*-expression confirmed that neurons with OPC contact had lower fluorescence intensity for *c-Fos* as compared to those without (Fig. 3e-f). Finally, we applied ATP (5 mM) and glutamate (1 mM) to stimulate the network in acute slices and compared the calcium responses between neurons with and without OPC contact. Neurons in contact with OPCs consistently exhibited significantly lower calcium responses under stimulating conditions (Fig. 3g), suggesting that OPCs modulate neuronal activity not only during spontaneous activity of the network but also during exogenous stimulation.

**Figure 3.**
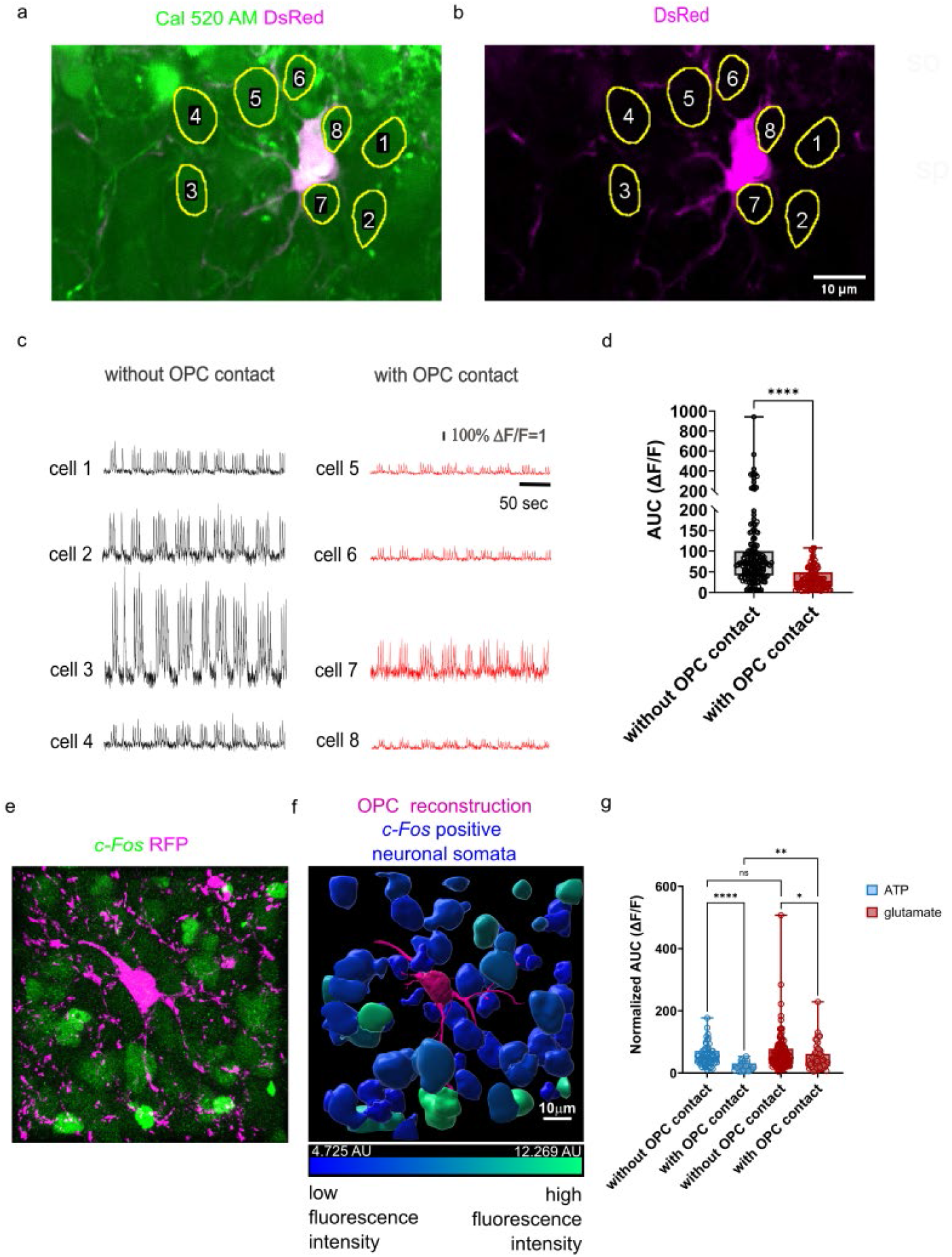
Neurons with contact to OPC processes show lower average calcium transients as compared to those without. a. Representative two-photon image of the hippocampal pyramidal layer in a neonate NG2DsRedBAC transgenic mouse. Neurons were bath-incubated with the Calcium Indicator Cal520 ®. Yellow circles mark neurons with or without nearby OPC contact. Scale bars: 10µm. b. Same field of view showing DsRed expression in OPCs and the spatial relationship with surrounding neurons. Scale bars: 10µm. c. Example traces of neurons recorded from the pyramidal layer of the hippocampus with and without OPC contact. d. Comparison of area under the curve (AUC) between neurons with or without OPC contact. Recording time=300 seconds, n=284 cells, 107 neurons had close contact to OPCs, 177 were without contact to OPCs. Shapiro-Wilk test was performed to assess normality (p<0.0001) and Mann-Whitney test was performed subsequently (p=<0.0001, two-tailed, sum of ranks in column A=30851; B=9619). e. Confocal image of acute slice after fixation and immunohistochemical labeling of red fluorescent protein (indicating expression of DsRed, shown in magenta) and *c-Fos* (green). Scale bars: 10µm. f. 3D reconstruction showing lower fluorescence intensity of *c-Fos* labeling (intensity code shown underneath the image) in neurons directly contacted by an OPC (blue) compared to distal neurons (green). Scale bars: 10µm. g. Comparison of AUC after application of ATP and glutamate in neurons with and without OPC contact. AUC was normalized to the AUC of Alexa594 as a measure for the amount of agonist applied. Recording time: 300 seconds. Kruskal–Wallis test with Dunn’s post hoc correction for multiple comparisons was performed (ATP: without OPC contact vs with OPC contact p<0.0001; glutamate: without OPC contact vs with OPC contact p=0.0227; ATP without OPC contact vs glutamate without OPC contact p>0.9999; ATP with OPC contact vs glutamate with OPC contact p=0.0013). ATP group: n = 89 cells; glutamate group: n = 214 cells.

In summary, we show that OPC processes are frequently localized around neuronal somata in different brain regions and age groups. We provide evidence that these contacts are established to detect and dynamically respond to changes in neuronal activity and can even be used to influence and regulate neuronal activity. Supporting the hypothesis that this is a common physiological process, we detected significantly lower fluorescence intensity of immunohistochemically detected expression of the immediate early gene *c-Fos* in neurons surrounded by OPC processes in adult mice, under physiological conditions and after inducing an increase in neuronal activity in the network through chemogenetic stimulation. We could expand our findings to acute slices from neonate mice, where we observed lower calcium signaling in neurons contacted by OPCs in the neonate hippocampus. Although the exact mechanism by which OPCs reduce neuronal activity remains unresolved, possible explanations include physical contact limiting the access of signals or solutes to the surface of these neurons, or active release of signaling molecules by OPCs leading to intracellular changes in the contacted neuron. The observation that OPCs frequently and across developmental stages approach the soma of neurons is new, and aligns with other recently published studies implying that OPCs exert additional functions independent of myelination ^1,2,4,15^. Interestingly, microglia have also been shown to form specialized somatic purinergic contacts with neurons ^16^. These contacts can modulate neuronal activity by suppressing neuronal activity ^17^ and exert a neuroprotective role during homeostasis that is more pronounced after brain injury ^16^. We add OPCs as novel modulators to be positioned at neuronal structures that allow for the detection and regulation of neuronal activity. Along these lines, we found microglia and OPCs to be co-localized at the AIS to a comparable extent (Extended Data Fig. 5).

Our results suggest that OPCs are substantially involved in regulating the activity of the neural network, specifically reducing the activity levels of neurons they surround, potentially exerting developmental, coordinating and neuroprotective functions.

## ACKNOWLEDGEMENTS

We thank Prof. Dr. Akiko Nishiyama (University of Connecticut, Storrs) for providing the tissue for the analysis of the juvenile Scn1a mutant mice and the adult DREADD mice. We thank Dr. Yury Kovalchuk for help with establishing the measurement of calcium transients in the NG2DsRed mice. We thank Prof. Dr. Matthew Rasband (University of Connecticut Health Center, Farmington) for the generous donation of the rabbit anti-β4-spectrin antibody. We thank Youfen Sun for maintaining the mouse colony at the Nishiyama lab. We thank Elizabeta Zirdum and Kerrin Schmidt for excellent technical assistance. We thank Dr. Chris O’Connell, Director of Advanced Light Microscopy Facility, for his assistance with the Leica SP8 confocal microscope, which was purchased with funds from an NIH Instrumentation Grant S10 OD016435 (PI: Akiko Nishiyama). We thank Dr. Olga Oleksiuk at the HIH CIN Imaging Cluster, Tübingen, for support with using the Leica SP8 confocal microscope.

The work was supported by the European Union’s Framework Program for Research and Innovation Horizon 2020(2014-2020) under the Marie Sklodowska-Curie Grant Agreement No. 845336 to FP. YS is supported by the China Scholarship Council (grant number 202309370022). YA was supported by the IBACS Summer Graduate Fellowship from the University of Connecticut Institute of Cognitive Sciences and the Kenneth and Paula Munson Family Fund for student support in Health Sciences Fellowship from the University of Connecticut Institute of Systems Genomics.

