## Supplementary material for "Oligodendrocyte precursor cells establish contacts with somata of active neurons": BIORXIV2025646001_supplementary information.pdf

**Extended Data Figures**

**Extended Data Figure 1. Distribution of OPCs in the CA1 region of the P14 hippocampus.**

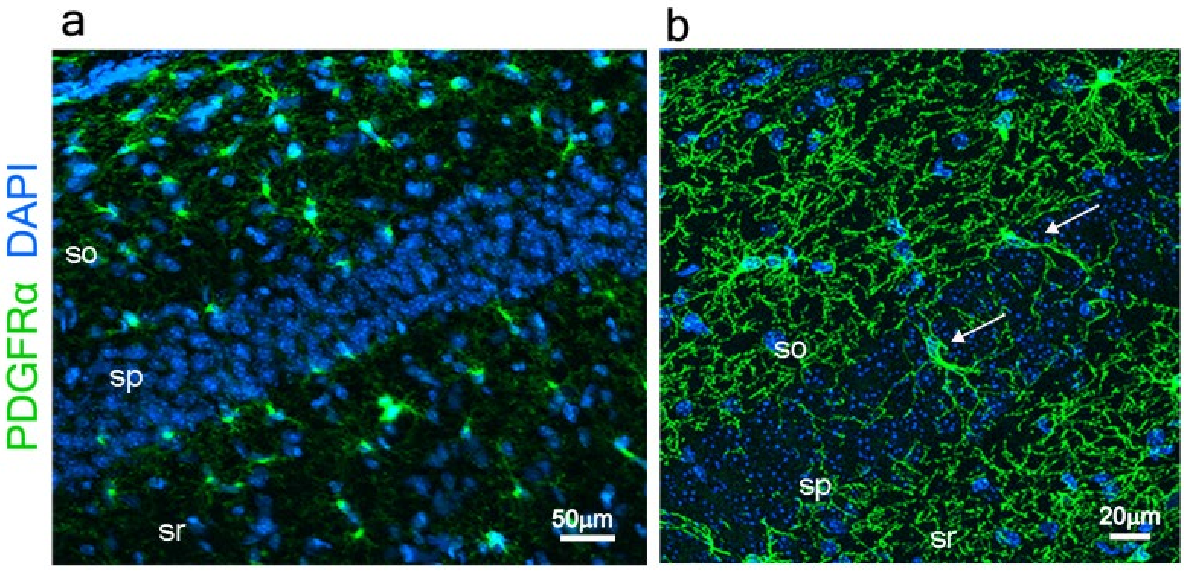


a. Representative epifluorescence image of the CA1 hippocampal region showing the distribution of OPCs (stained with anti-PDGFRα antibody in green) in stratum oriens (so), stratum pyramidale (sp), and stratum radiatum (sr). DAPI is shown in blue for better orientation. Scale bar: 50µm.

b. Zoomed in confocal images showing OPCs (stained with anti-PDGFRα antibody in green) inserting processes into the stratum pyramidale (white arrows). Scale bar: 20µm.

**Extended Data Figure 2. Juvenile model for increased neuronal activity**

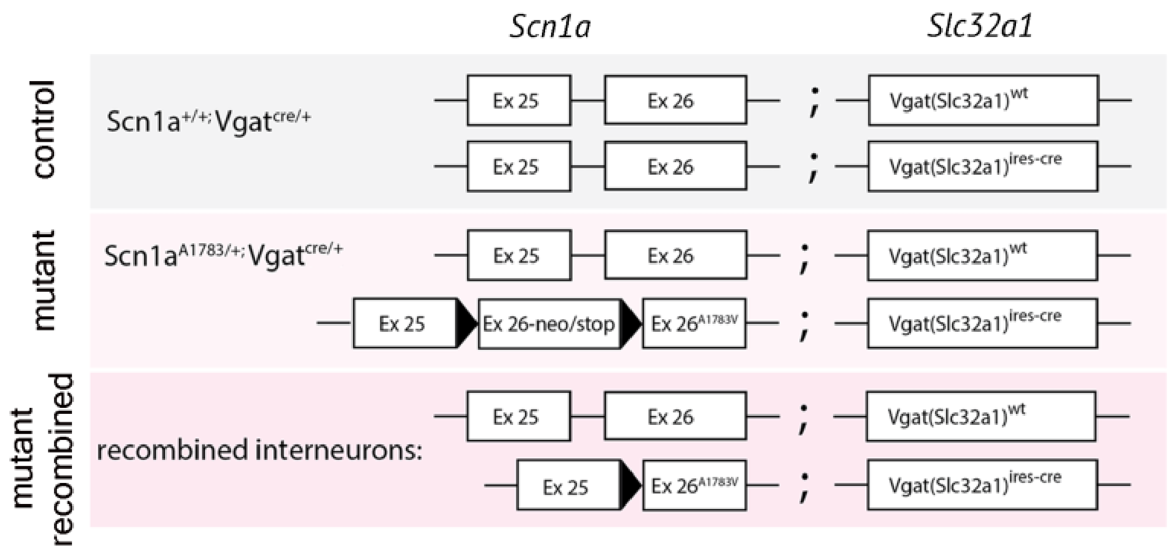


The Scn1a mutant mice express the loss-of-function Scn1a-A1783V mutation encoding the voltage-dependent Nav1.1 sodium channel conditionally in GABAergic inhibitory interneurons (Kuo, Cleary et al. 2019, Ricobaraza, Mora-Jimenez et al. 2019), leading to hyperactivity of principal neurons due to reduced inhibitory input.

A schematic showing *Scn1a* and *Slc32a1 (Vgat)* alleles in Scn1a control (gray) and Scn1a mutant (pink) mice. Ex 25 and Ex 26 represent exons of the *Scn1a* gene. Genotype in the middle (light pink) shows before recombination. Bottom in darker pink shows genotype in recombined interneurons expressing Cre. Black triangles, loxP sites. The mutant mice developed normally during the first two weeks after birth. At postnatal day 14 (P14) occasional seizures started to appear, and the mice died after P15 due to respiratory problems, as previously described (Kuo et al., 2019, Ricobaraza et al., 2019).

**Extended Data Figure 3. Density of OPCs in the hippocampus is not changed in Scn1a mutant mice as compared to controls at P14.**

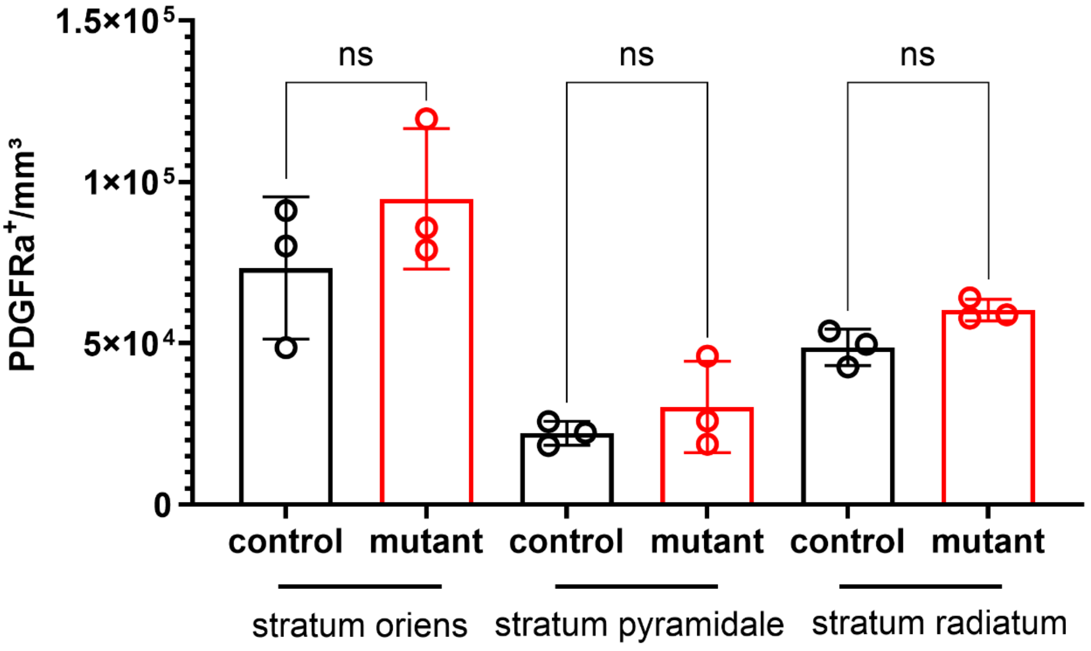


Density of OPCs in stratum oriens, stratum pyramidale and stratum radiatum of the hippocampus. Ordinary one-way ANOVA was performed with Tukey's multiple comparisons test. Treatment (between columns) F (5,12) =10.89, P=0.0004. n=3 mice.

**Extended Data Figure 4. OPCs in Scn1a mutant mice do not have more processes compared with their age-matched control littermates in the hippocampal CA1 at P14.**

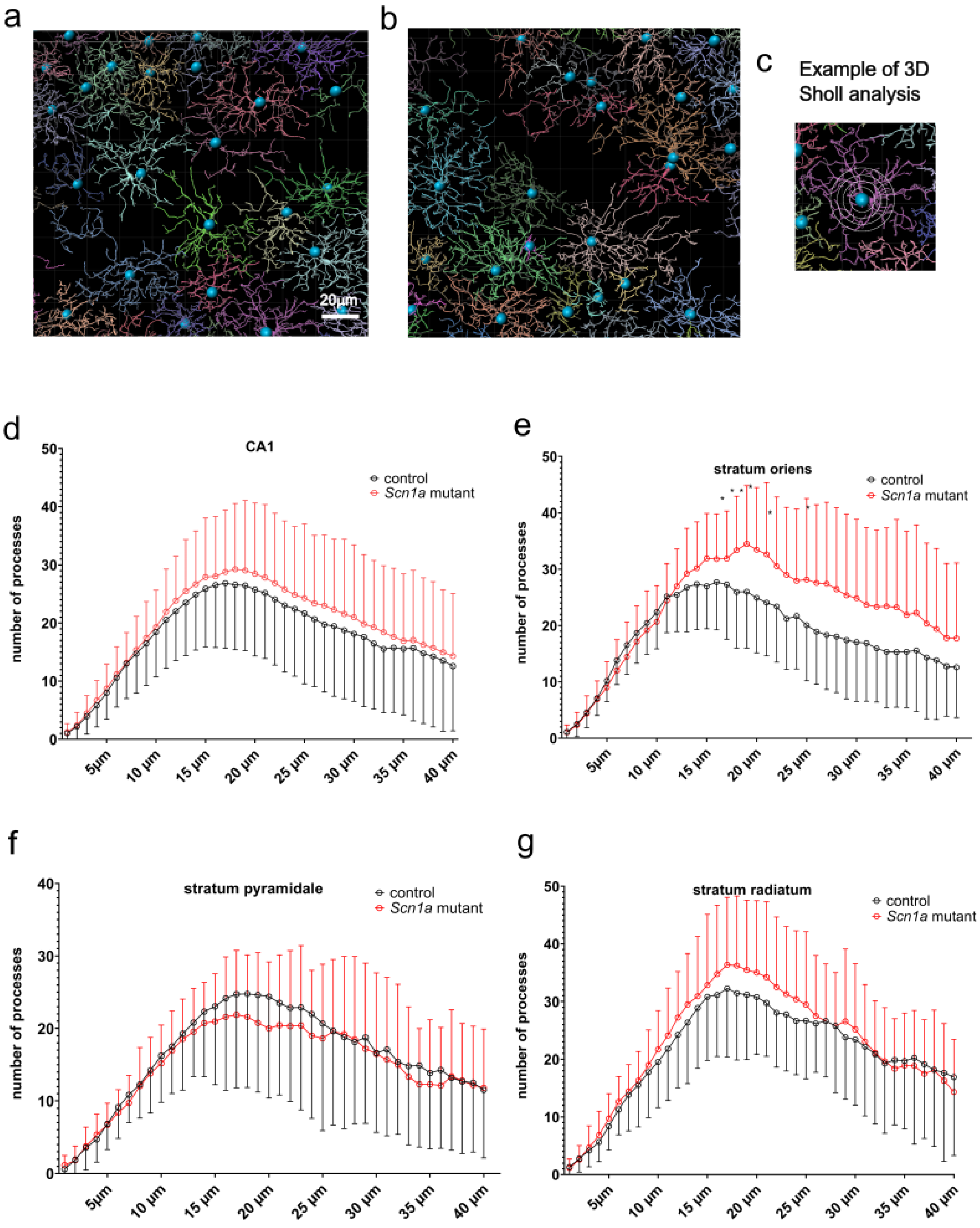

Automated filament reconstructions and Sholl analysis.

a. Example of 3D reconstructions of OPCs for Sholl analysis in control mice generated in Imaris using the filament tracer module. Scale bar: 20μm.
b. Example of 3D reconstructions of OPCs for Sholl analysis in SCn1a mutant mice generated in Imaris using the filament tracer module. Scale bar: 20μm.
c. Example of Sholl speres with different radii around each OPC reconstructed.
d. Sholl analysis from 3D reconstructions of the number of OPC processes at indicated distances from soma center in the CA1 of Scn1a mutant mice and controls (n=136 OPCs per group). 2-way ANOVA and Sidak's multiple comparison test were applied. Row Factor x Column Factor = 0.3564.
e. Sholl analysis from 3D reconstructions of the number of OPC processes at indicated distances from soma center in the stratum oriens of Scn1a mutant mice (n=41 OPCs) and controls (n=46 OPCs). 2-way ANOVA and Sidak's multiple comparison test were applied. $p \leq 0.05$ for the following distances: 18μm=0.0271, 19μm=0.0084, 20μm=0.0119, 21μm=0.0280, 23μm=0.0428 and 27μm=0.0440. Row Factor x Column Factor = 0.0008.
f. Sholl analysis from 3D reconstructions of the number of OPC processes at indicated distances from soma center in the stratum pyramidale of Scn1a mutant mice (n=25 OPCs) and controls (n=21 OPCs). 2-way ANOVA and Sidak's multiple comparison test were applied. Row Factor x Column Factor = 0.7157.
g. Sholl analysis from 3D reconstructions of the number of OPC processes at indicated distances from soma center in the stratum radiatum of Scn1a mutant mice (n=30 mice) and controls (n=27 OPCs). 2-way ANOVA and Sidak's multiple comparison test were applied. Row Factor x Column Factor = 0.3845.

**Extended Figure 5: Microglia and NG2 cells colocalize with the AIS in similar patterns and at similar frequency in the adult mouse.**

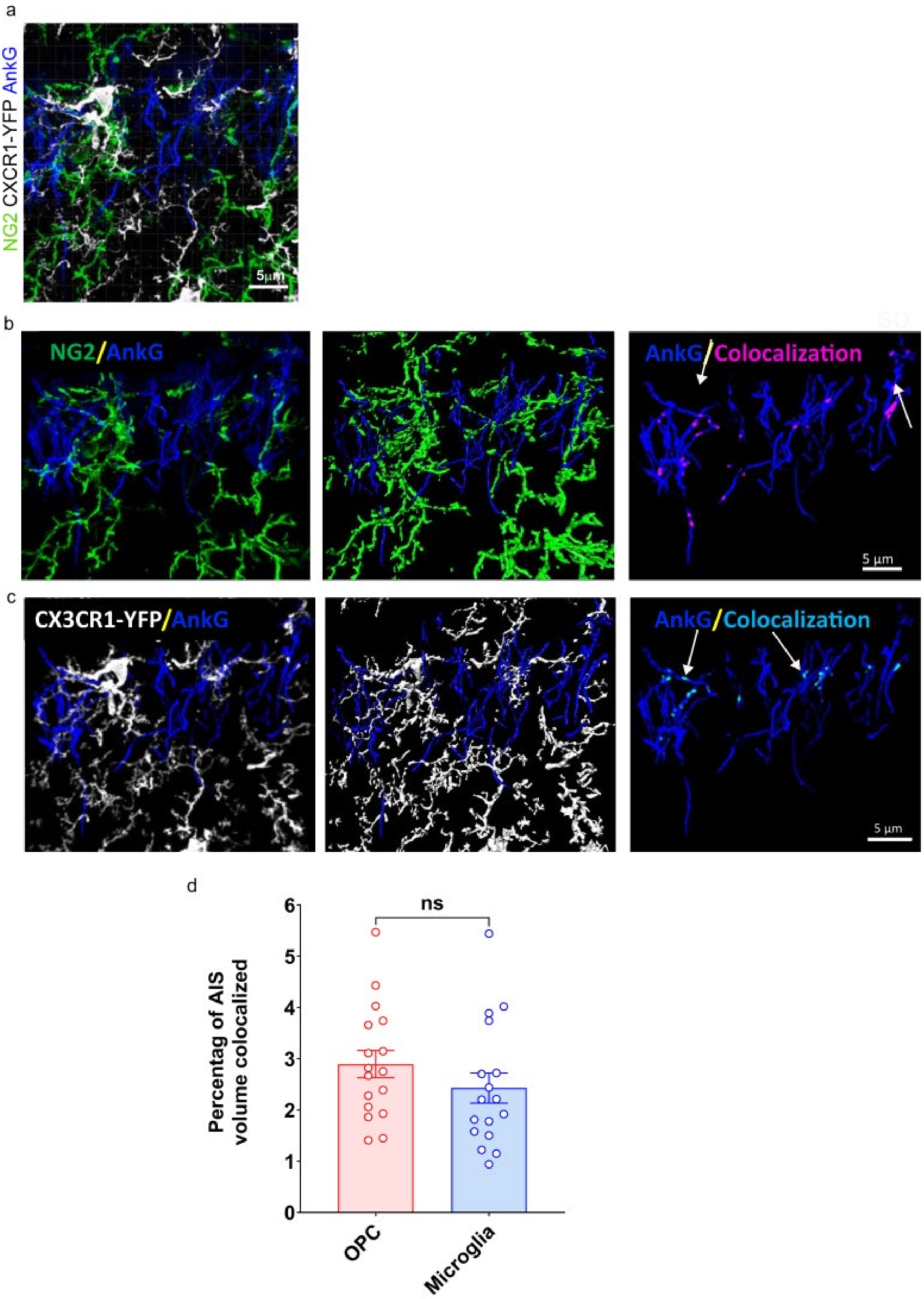

a. Imaris 3D reconstruction that was used to quantify colocalization. OPC processes were stained with anti-NG2 antibody (shown in green), the axon initial segment (AIS) was stained with anti-Ankyrin G (shown in blue) and microglial processes are shown in white (CXCR1-YFP mouse). Scale bar: 5μm.

b. OPC processes (stained with anti-NG2 antibody, shown in green) colocalize with the AIS (stained with anti-AnkG antibody, shown in blue) in the hippocampal CA1 region. Imaris 3D isosurface rendering (middle panel) was used to mask background in the immunostained image (left panel). Colocalization is indicated in the right panel in magenta (white arrows). Scale bar: 5μm.

c. Microglial processes (CXCR1-YFP mouse, shown in white) colocalize with the AIS (stained with anti-AnkG antibody, shown in blue). Imaris 3D isosurface rendering (middle panel) was used to mask background in the immunostained image (left panel). Colocalization is indicated in the right panel in cyan (white arrows). Scale bar: 5μm.

d. Quantification of the percentage of AIS volume that was covered by microglial (~2.5%) and OPC (~3%) processes. A paired t-test indicated no significant difference (p=0.2497; n=17 images from 3 mice).
